# A conserved graft formation process in *Picea abies* and *Arabidopsis thaliana* identifies the PAT gene family as central regulators of wound healing

**DOI:** 10.1101/2023.07.30.551201

**Authors:** Ming Feng, Ai Zhang, Van Nguyen, Anchal Bisht, Curt Almqvist, Lieven De Veylder, Annelie Carlsbecker, Charles W. Melnyk

## Abstract

The widespread use of plant grafting has enabled eudicots and gymnosperms to join with closely related species and grow as one. Gymnosperms have dominated forests for over 200 million years and despite their economic and ecological relevance, we know little about how they graft. Here, we developed a micrografting method in conifers using young tissues that allowed efficient grafting between closely related species and distantly related genera. Conifer graft junctions rapidly connected vasculature and differentially expressed thousands of genes including auxin and cell wall-related genes. By comparing these genes to those induced during *Arabidopsis thaliana* graft formation, we found a common activation of cambium, cell division, phloem and xylem-related genes. A gene regulatory network analysis in *Picea abies* (Norway spruce) predicted that *PHYTOCHROME A SIGNAL TRANSDUCTION 1* (*PAT1*) acted as a core regulator of graft healing. This gene was strongly upregulation during both *P. abies* and *Arabidopsis* grafting, and *Arabidopsis* mutants lacking *PAT*-genes failed to attach tissues or successfully graft. Complementing *Arabidopsis* PAT mutants with the *P. abies PAT1* homolog rescued tissue attachment and enhance callus formation. Together, our data demonstrate an ability for young tissues to facilitate grafting with distantly related species and identifies the PAT gene family as conserved regulators of graft healing and tissue regeneration.

## Introduction

The cutting and joining of different plants during the process of grafting has been practiced for millennia to combine the best properties of two plants. Grafting likely originated as a means for vegetative propagation but now is commonly used to improve stress tolerance, to retain varietal characteristics and to enhance yields. Today, the majority of commercially grafted plants are eudicots but gymnosperms too are grafted to propagate desirable varieties for forestry breeding programs and in horticulture ^1–4^. Gymnosperms evolved approximately 200 million years before angiosperms, and today, dominate many forest environments and have important economic and ecological consequences ^4, 5^. Despite their widespread prevalence and their ability to be grafted, we have little understanding of how such a process might function in gymnosperms and its relationship to grafting in angiosperms. In conifers, our ability to successfully graft is limited by various factors including grafting techniques, grafting season, pathogen contamination and the relatedness of species ^6–8^. Closely related conifer or eudicot species from the same genus normally successfully graft whereas combinations from different genera often fail, a phenomenon known as graft incompatibility that limits grafting success^9,10^. The mechanistic basis for graft incompatibility remains unclear but might be due to structural weakness, metabolic imbalances or the activation of defense responses ^11–13^. However, not all distantly related grafts fail and inter-genus grafts within the cactus and Solanaceae families are possible ^12^. Recently, inter-family grafts were made with *Petunia hybrida* or *Nicotiana benthamiana* where a cell-wall related β*-1,4- glucanase* gene is important to promote graft attachment ^14, 15^. Protocols to successfully graft monocots have recently been established using embryonic tissues which allow both inter- and intra-species grafts ^16^. Grafting with such small tissues, a process known as micrografting, is increasingly being used as a tool to improve grafting efficiency and also to characterize the process of grafting itself ^16–19^.

Work in tomato, *Sedum* and *Arabidopsis* has revealed a dynamic healing process at the graft junction ^17, 20, 21^. After cutting, cells expand and divide to adhere tissues and fill the wound. Cell wall components, including pectins, are secreted and the expression of cell wall-related genes such as β*-1,4-glucanase* plays an important role in the early stages of graft attachment ^14, 22^. Cell divisions lead to the formation of callus, a stem-cell like tissue, at the cut ends that helps seal the wound. In the final stages of graft formation, the callus and surrounding tissues are differentiated to functional phloem and xylem tissues and outer cell layers to resume vascular transport and reform protective barriers. During grafting, thousands of genes are differentially expressed including early activating transcription factors such as *ETHYLENE RESPONSE FACTORs* (*ERFs*), *DNA binding with one finger* (*DOF*) transcription factors and NAC domain-containing proteins (*ANACs*) ^23, 24^. These factors play important roles during grafting to promote tissue adhesion, callus formation and vascular differentiation ^23–25^. In addition to its role in grafting, *ERF115* also controls the replenishment of cells after wounding in the *Arabidopsis* root meristem. The regenerative ability of *ERF115* is enhanced by its interacting partner *PHYTOCHROME A SIGNAL TRANSDUCTION1 (PAT1). PAT1*, along with two other GRAS transcription factors, *SCARECROW-LIKE5* (*SCL5*) and *SCL21*, are important for root tip regeneration and cell death recovery ^26, 27^. *ERF115* also plays an important role during wounding to enhance auxin sensitivity by activating *AUXIN RESPONSE FACTOR5* (*ARF5)* ^28^. Auxin is important for graft formation since blocking auxin transport inhibits the ability of *Arabidopsis* and rice grafts to heal, whereas reducing auxin response below the junction inhibits *Arabidopsis* graft healing ^16, 17, 29^. Other early activators during grafting include *WUSCHEL-RELATED HOMEOBOX13* (*WOX13*) and *WOUND INDUCED DEDIFFERENTIATION1 (WIND1)* that are important for callus formation at the site of cutting ^30, 31^. Thus, work in *Arabidopsis*, rice and *Nicotiana* has identified a number of factors that are activated early and contribute to attachment, callus formation and vascular differentiation during grafting.

Here we investigated the process of graft formation in several widespread and commercially relevant conifer species. We developed an efficient and practical grafting method using young conifer plants that allowed graft junctions to rapidly heal and permitted several inter-species and inter-genus graft combinations to form successfully. We used this method to characterize graft healing and discovered a common graft formation pathway in *Picea abies* (Norway spruce) and *Arabidopsis* that involved cell division, vascular differentiation and the upregulation of cell wall and auxin related genes. We additionally identified that *PAT1* upregulation is common in *Arabidopsis* and *P. abies* grafting and that this gene appears to have a conserved role in wound healing between gymnosperms and eudicots.

## Results

### A new method for conifer grafting

Previous conifer grafting methods were limited by various factors including techniques, grafting season and contamination ^6, 7^. To improve graft formation rates and the ease of grafting, we developed a micrografting method using 10-12 days old *Picea* (spruce) and *Pinus* (pine) seedlings. Plants were excised in the hypocotyl region, and scions and rootstocks from different plants were attached tightly together using a silicon collar (Fig. 1a, b). With practice, 50 grafts per hour could be made with >90% success rates (Table 1). To monitor the dynamics of graft healing, we grafted *P. abies* to *P. abies* or *Pinus contorta* (Lodgepole pine) to *P. contorta* and treated the scion and rootstock with carboxyfluorescein diacetate (CFDA), a dye used for testing vascular connectivity in grafted *Arabidopsis* and rice ^16, 17^. 10 days after grafting (DAG), we treated plants with CFDA in the scion and observed nearly half of plants could transport the dye to the roots, consistent with resumption of shoot-to-root transport through the phloem (Fig. 1c, d). Similarly, CFDA application to the rootstock at 10 DAG showed movement to the scion in nearly half of plants consistent with resumption of root-to-shoot transport through the xylem (Fig. 1c, d). By 20-25 DAG, nearly all individuals showed transport dynamics consistent with phloem and xylem connectivity. As a second test of vascular reconnection, we stained hand sections from the graft junction with basic fuchsin to assess the presence of xylem associated-lignin. Successful grafts showed xylem connections across the junction, whereas unconnected plants showed little xylem staining and only callus formation at the junction (Fig. 1e, Extended Data Fig. 1). Two months after grafting, the grafted spruce and pine showed normal growth, well healed junctions and survival rates of 90∼100% (Fig. 1b; Supplementary Table 1). Thus, our technique was an efficient and practical method for grafting young conifers that allowed the graft junction to rapidly form xylem and phloem connections after grafting.

**Fig. 1.**
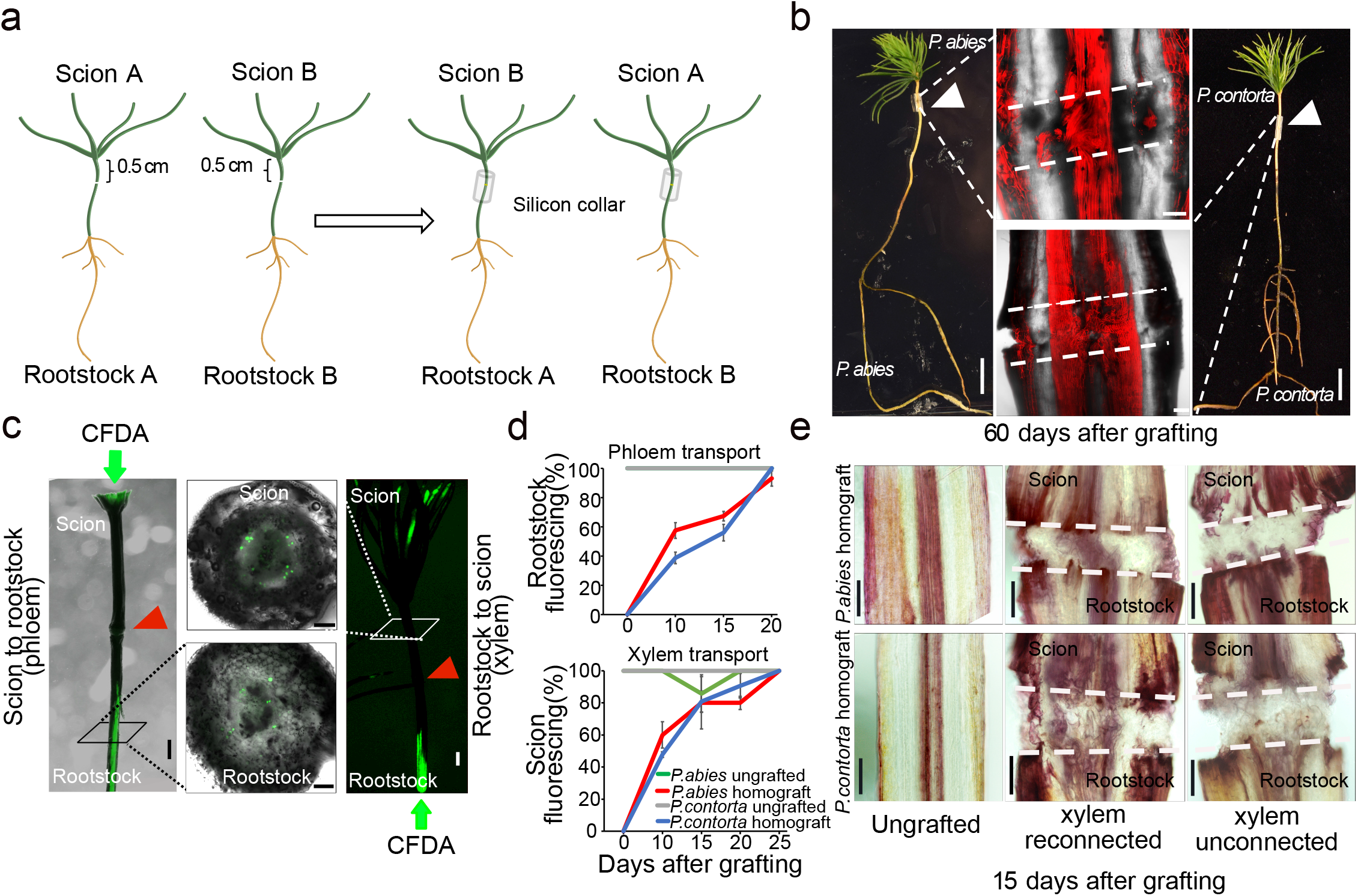
Micrografting dynamics in conifers. a, Cartons showing the micrografting method used in conifers. 10-12 days old scions and rootstocks from different plants are joined with the help of a silicon collar to grow as a new plant. b, Homografted *Picea abies* (left) and *Pinus contorta* (right) 60 days after grafting. Scale bars, 1cm. White triangles indicate the graft junction. Middle panels show confocal images of the vascular anatomy at the graft junction. Scale bars, 100 µm. c, Phloem and xylem transport assays in conifers involving CFDA application to the scion (phloem) or rootstock (xylem) monitored the appearance of fluorescence in rootstock or scion, respectively, consistent with phloem or xylem transport. Scale bars, 1mm. Hand-section stems above or below the graft junction confirmed vascular transport. Scale bars, 100 µm. d, Phloem and xylem reconnection rates in *Picea* and *Pinus* grafts. The mean from three biological replicates with 11-27 plants per time point per experiment is shown (± s.d.). e, Xylem staining 15 days after grafting in *Picea* or *Pinus*. Xylem was stained with basic fuchsin and showed reconnection in successful grafts. Scale bars, 20 µm.

**Table 1.**
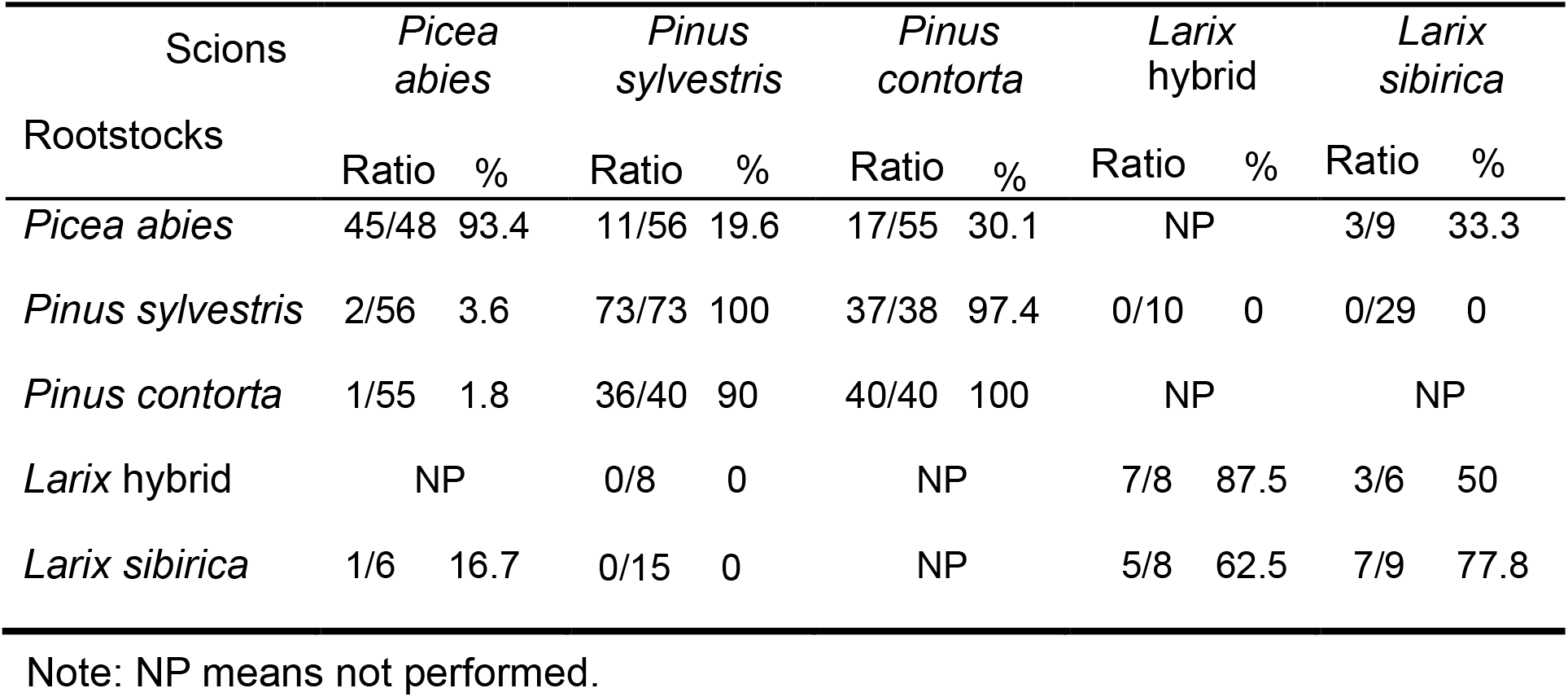
Graft success rate in different conifer combinations after 2.5 years of growth.

### Micrografting improves heterograft compatibility

Previous studies showed that grafting success decreased and incompatibility increased as conifer species became more distantly related ^3, 9^, therefore, we tested our micrografting method to determine whether we observed similar results. Since the Pinaceae family shows the closest relatedness between *Picea* and *Pinus* (Fig. 2a)^32^, we first tested grafting between *Picea* and *Pinus* species (heterografting). Two months after grafting, species grafted to themselves (homografts) or different species (heterografts) had high survival rates varying between 70-100% depending on the genotype (Supplementary Table 1). We assessed xylem connectivity in the *P. contorta*/*P. abies* heterograft (scion/rootstock notation) and found xylem connectivity was similar to homografted plants (Fig. 2b). However, incompatibility in woody species can develop several years after grafting ^33^, we therefore moved plants to soil for long term observation. 2.5 years after grafting, the survival rates of homografted plants remained high (90%∼100%) but heterografted *Picea*-*Pinus* combinations showed lower survival rates (Table 1). *P. abies* scions grafted to *Pinus sylvestris* (Scots pine) or *P. contorta* rootstocks had low survival rates (3.6% and 1.8% viable, respectively), but *P. abies* rootstocks performed better when grafted to *P. sylvestris* or *P. contorta* scions (19.6% and 30.1% viable, respectively) (Table 1). Successful grafts displayed good growth, though some heterografted combinations were shorter and showed swelling at the graft junction. Heterografted plants without swelling had similar heights to the intact plants (Fig. 2c-f, Supplementary Table 2). *P. abies* rootstocks also appeared to reduce the needle length of the *Pinus* scions (Fig. 2c, e). Next, we tested two species from the Pinaceae genus *Larix* in heterografts (Extended Data Fig. 2). *Larix sibirica*/*P. sylvestris* combinations did not survive whereas *L. sibirica*/*P. abies* had three of nine plants growing well but with swelling at the graft junction (Table 1, Fig. 2). One of six *P. abies*/*L. siberica* survived, but it showed poor growth (Table 1, Extended Data Fig. 2). Our results suggested that micrografting allowed several inter-species and inter-genus grafts to successfully form and that *P. abies* rootstocks appear to inable long term grafting success with divergent scions.

**Fig. 2.**
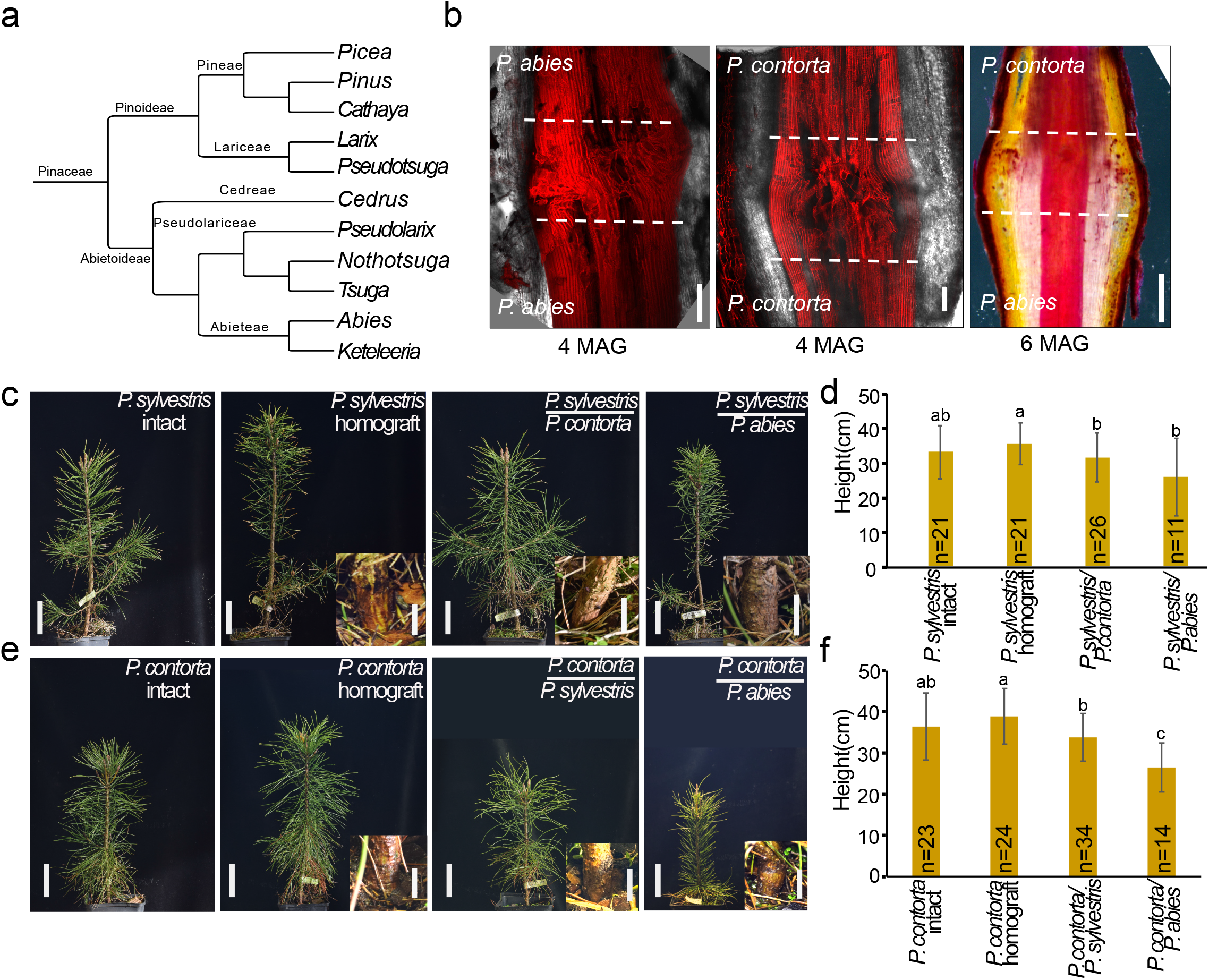
Micrografting enables inter-genus grafting. a, Phylogenetic tree showing the intergeneric relationships of Pinaceae based on Ran et al. 2018. b, Junction anatomy of homografted or heterografted *Pinus contorta* and *Picea abies* 4 or 6 months after grafting (MAG). Scale bars, 100 µm for images at 4 MAG, 1 mm for image at 6 MAG. (c, e), Representative images of homografted or heterografted *Pinus sylvestris*, *Pinus contorta* and *Picea abies* 2.5 years after grafting. Intact (non-grafted) *Pinus sylvestris* and *Pinus contorta* were used as controls. Scale bars, 10 cm. Inserts show the graft junctions. Scale bars, 1cm. (d, f), Height of intact and 2.5 years old grafted *Pinus sylvestris*, *Pinus contorta* and *Picea abies* (n=11-34 per combination).

### Grafting activates vascular and cell division-associated genes

To understand the process behind conifer grafting and to identify the genes differentially expressed, we generated RNAseq libraries from both ungrafted (intact) and grafted *P. abies* above and below the graft junction at 0, 1, 3, 7,14 and 28 DAG (Fig. 3a). A Principal Component Analysis (PCA) showed samples largely clustered by tissues type and time point with a close correlation between scion and rootstock samples (Fig. 3b). Intact and grafted samples had similar numbers of expressed genes yet there was an increase in differential expression in grafted samples particularly at 1 and 3 days after grafting compared to intact controls (Extended Data Fig. 3a-d, Supplementary Table 3). To analyse common patterns of gene expression in grafted tissues, we used Mfuzz to group differentially expressed genes into 12 clusters (Fig. 3c-e, Extended Data Fig. 3e, f). These including genes upregulation in both scion and rootstock (cluster 5,6,7,11,12), scion-specific upregulation (cluster 10), rootstock-specific upregulation (1,2,9) or downregulation in scion and rootstock (cluster 3,4,8). A gene ontology (GO) analysis on the clusters revealed enrichment of wounding and defense-related processes in cluster 2, enrichment of cell cycle-related processes in cluster 5, and enrichment in cell wall and xylem-related processes in cluster 10 (Supplementary Table 4). Within these clusters, we searched for homologs of previously described grafting-related genes ^23, 30, 34^ to better understand how conifers graft. Cluster 6 contained early activating genes in the scion and rootstock including a wounding-related *PaWOX13-like* gene ^30^(Fig. 3d, g). Cluster 10 contained early activating scion-specific genes including a wounding-related *PaWIND1-like* gene (Extended Data Fig. 3c). Cluster 5 contained genes that slightly later increased in both rootstock and scion including cell cycle related genes such as *PaCDKB2;2-like* (Fig. 3c, f) (Supplementary Table 4). A phloem-related *PaAPL-like*, xylem-related *PaPRX66-like* and *PaVND4-like* genes had intermediate and late activation dynamics (Fig. 3e, h, Extended Data Fig. 3g, h). To compare gene expression profiles in grafted conifers and eudicots, we analysed the grafting transcriptomics dataset from *Arabidopsis* ^23^ and found the expression of several *P. abies* gene homologs had similar expression patterns in Arabidopsis (Figure 3f-h, Extended Data Fig. 3e). Thus, there appeared to be consistent activation of homologous genes during *Arabidopsis* and *P. abies* grafting suggesting a conserved grafting process involving wound response, followed by cell division, and phloem and xylem differentiation.

**Fig. 3.**
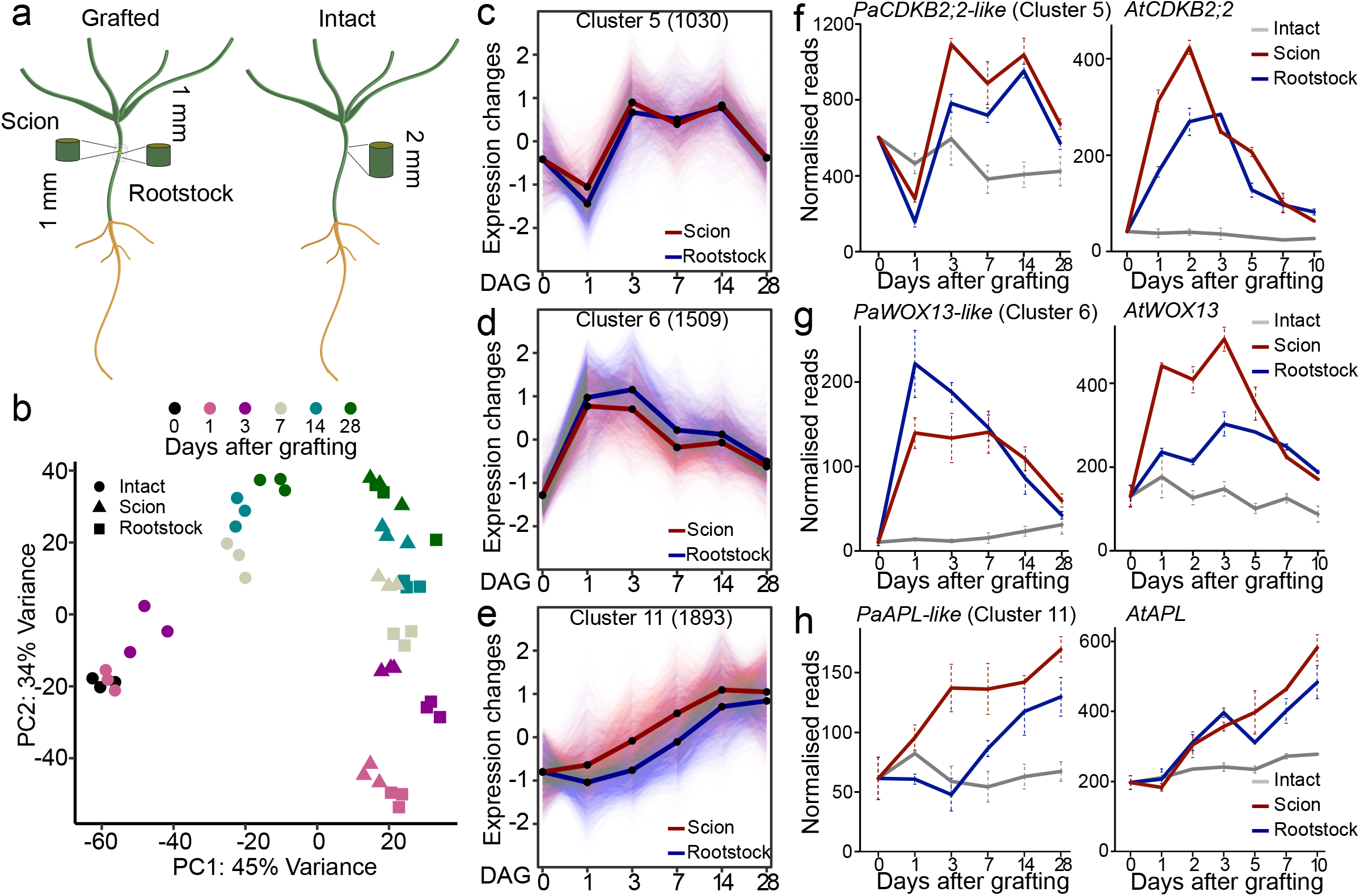
Transcriptome dynamics during *Picea abies* graft healing. a, Schematic diagram showing where *Picea abies* tissues 1 mm above or 1mm below the graft junction were harvested as scion and rootstock material, respectively, for transcriptome sequencing. b, Principal component analysis of the gene expression data from *Picea abies* graft healing transcriptomes. Colors indicate the different days after grafting (DAG), shapes indicate different tissues. (c, d and e), Clustering analysis of transcriptional dynamics during graft healing. Lines indicate the average of differentially expressed genes in scion or rootstock. Dots indicate days after grafting. The number in the brackets represents the number of genes in the cluster. (f, g and h). Expression profiles for select *Picea abies* genes belonging to clusters in (c, d, e). Cluster number is indicated. *Arabidopsis* homolog expression data are plotted and taken from published transcriptome data ^20^.

### Auxin responses increase and correlate with cell wall-related gene expression

Auxin and cytokinin play important roles during vascular formation ^35, 36^. We explored whether they were relevant for *P. abies* graft formation. Auxin and cytokinin-responsive genes in the hypocotyl are not well described in *P. abies* so we first treated two weeks old seedlings with a synthetic cytokinin 6-benzylaminopurine (BAP), auxin (IAA) or BAP plus IAA for 0, 5, 10 or 15 days (Fig. 4a). We harvested 8-10 mm of treated *P. abies* hypocotyl tissues and used these for RNAseq analyses. A PCA grouped samples together largely based on hormone treatment rather than time point (Extended Data Fig. 4a). Looking at the individual time points, we found several thousand genes were auxin responsive or responded to both hormones, while a slightly lower number responded to cytokinin (Fig. 4b-d, Extended Data Fig. 4b, Supplementary Table 5). To focus on genes specifically induced by auxin or cytokinin, we looked for genes induced at all three time points but induced only by the presence of cytokinin or auxin alone. We found auxin induced 2598 genes and repressed 2013 genes which we defined as auxin responsive genes. Cytokinin induced 710 genes and repressed 978 genes which we defined as cytokinin responsive genes (Fig. 4c, d). The average expression of these auxin responsive genes showed similar values in scions and rootstocks but cytokinin responsive genes showed a slightly higher expression in the scion at early time points (Extended Data Fig. 4c, d). Auxin induced genes were enriched in the scion while auxin repressed genes were enriched in the rootstock from 3 DAG onwards (Fig. 4e). Cytokinin induced genes showed little enrichment in the scion or rootstock but cytokinin repressed genes showed enrichment at later time points particularly in the rootstock (Fig. 4f).

**Fig. 4.**
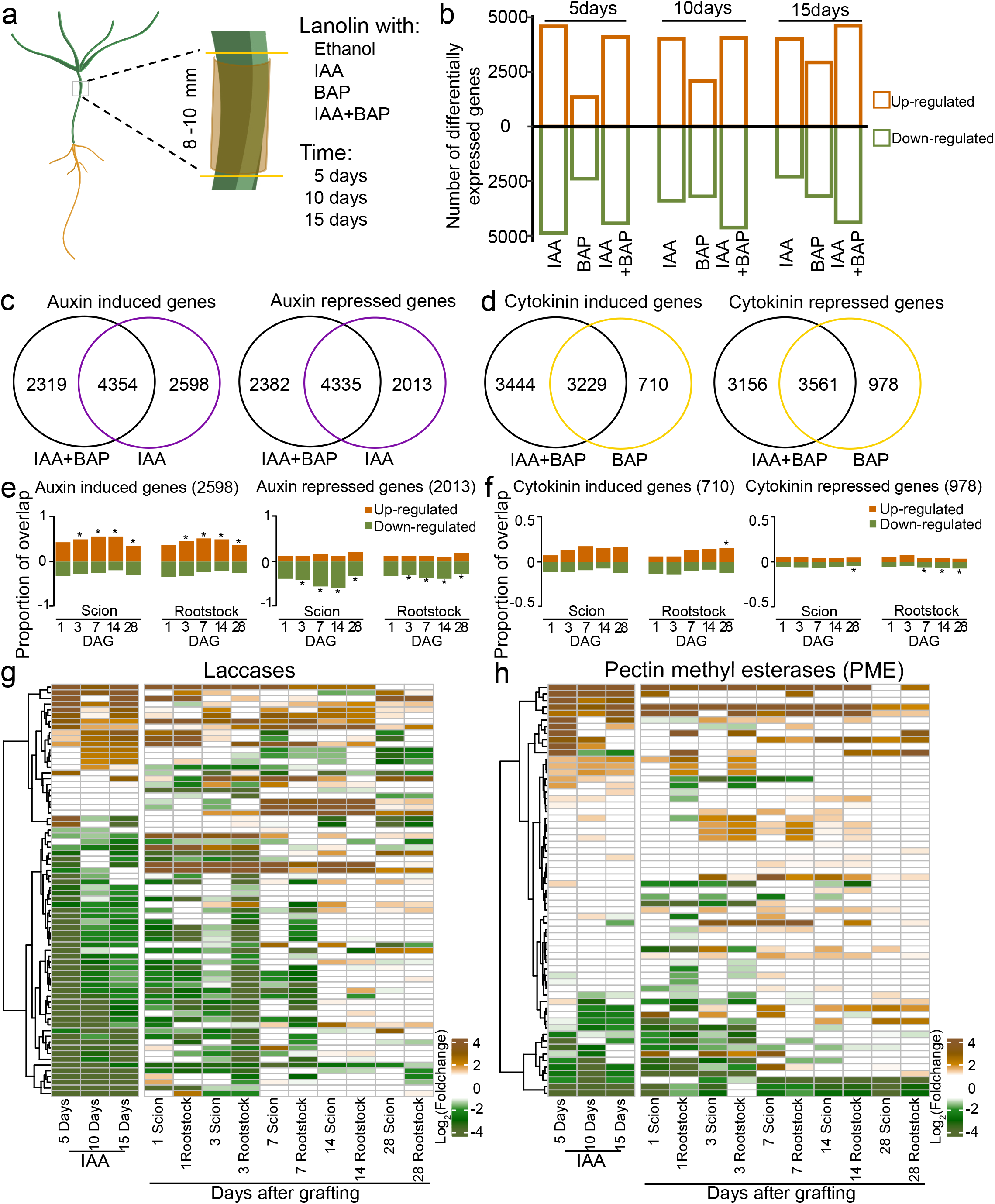
Auxin, cytokinin and cell wall related genes expression during grafting healing. a, Schematic diagram showing the regions harvested for the hormone transcriptomes in *Picea abies*. 50 mM auxin (IAA), 135µM cytokinin (BAP) or auxin plus cytokinin (IAA+BAP) was applied and tissues harvested 0 days, 5 days, 10 days and 15 days after treatment. b, The distribution of differentially expressed genes responding to hormone treatments. (c, d), Venn diagrams showing the auxin and cytokinin responsive genes differentially expressed by all three timepoints in the various treatments. (e, f), An overlap analysis between genes responsive to hormones (c and d) and genes differentially expressed during *Picea abies* grafting (Extended Data Fig.3c). Asterisks indicate statistically significant overlapped between hormone response and graft healing. \**p*<0.05. (g,h) Heatmap showing the fold changes of putative laccase genes (g) or putative pectin methyl esterases (PME)(h) in graft healing or auxin datasets.

Auxin response and cell wall modifications are important for successful graft formation ^14, 24, 37^ so we assessed whether auxin might affect cell wall related gene expression. Many putative *laccase*, *pectin methyl esterase* (PME), *beta-1-4-glucanase* and *pectate/pectin lyase* genes showed differential expression both after exogenous auxin treatment and after grafting including *PaLAC1*, *PaPME5-like*, *PaKOR1-like* and *PaPectin lycase-like* (Fig. 4g and h, Extended Data Fig. 4e-k). In particular, there was a substantial overlap between laccases and PMEs affected by both auxin treatment and grafting (Fig. 4e, f). These results demonstrated that grafting induced an auxin response at the junction and this correlated with the activation of cell-wall related genes at the graft junction.

### A conserved PAT1 gene family promotes graft healing

To further explore the transcriptional regulation of *P. abies* graft formation, we identified differentially expressed transcription factors (TFs) and mapped their abundance according to transcription factor gene families (Extended Data Fig. 5a). We performed a weighted correlation network analysis (WGCNA) and clustered these transcription factors according to their expression patterns in the grafting transcriptomes and defined 7 modules (Extended Data Fig. 5b, c). One module, represented in yellow (Extended Data Fig. 5c), showed differential expression specifically in scions and rootstocks but not in intact plants. Using the expression patterns of the genes in the yellow module, we generated a gene regulatory network (Fig. 5a, b). Most transcription factors were upregulated during grafting and in particular five, *PaPAT1-like*, *PaWIP4-like*, *PaMYB4-like*, *PaLRP1-like* and *PaMYB123-like*, were highly upregulated and appeared to act as hubs of the regulatory network (Fig. 5a, b, c, Extended Data Fig. 5d-g). We then used the regulatory network to test whether homologous genes were induced in *Arabidopsis*. We found that *Arabidopsis LRP1*, *MYB4* and *PAT1* were all induced during *Arabidopsis* grafting suggesting a broadly conserved regulatory response between *Arabidopsis* and *P. abies* (Fig. 5c, Extended Data Fig. 5d, e, g). Since the *PAT1* gene family promotes root tip regeneration ^26, 27^, we focused on this family and tested an *Arabidopsis PAT1* overexpression line (*AtPAT1OE*), *pat1scl5* mutant and *pat1scl5scl2* mutant in callus formation assays since callus is relevant for graft healing ^12, 38, 39^. The *AtPAT1OE* line showed increased callus formation at wounding sites of petioles, while *pat1scl5* and *pat1scl5scl2* showed strong impairment in callus formation (Fig. 5d, e). We also tested *Arabidopsis* graft attachment rates and found *pat1scl5*, *scl5scl21*, *pat1scl21* and *pat1scl5scl21* all reduced attachment rates (Fig. 5f). In CFDA-mediated phloem reconnection assays, the *PAT1OE* line showed no changes but the single, double and triple *PAT1*-related mutants all showed moderate to strong inhibition of phloem reconnection (Fig. 5g). ERF115 can interact with PAT1 family genes and *AtERF115*-*AtPAT1OE* co-overexpression induced callus formation (Extended Data Fig. 7)^26, 27^. We tested attachment and phloem reconnection rates in *erf115*, *erf115pat1* and *erf115scl5* mutants. The *erf115scl5* mutant failed to attach whereas the *erf115*, *erf115pat1* and *erf115scl5* mutants all inhibited phloem reconnection (Extended Data Fig. 7).

**Fig. 5.**
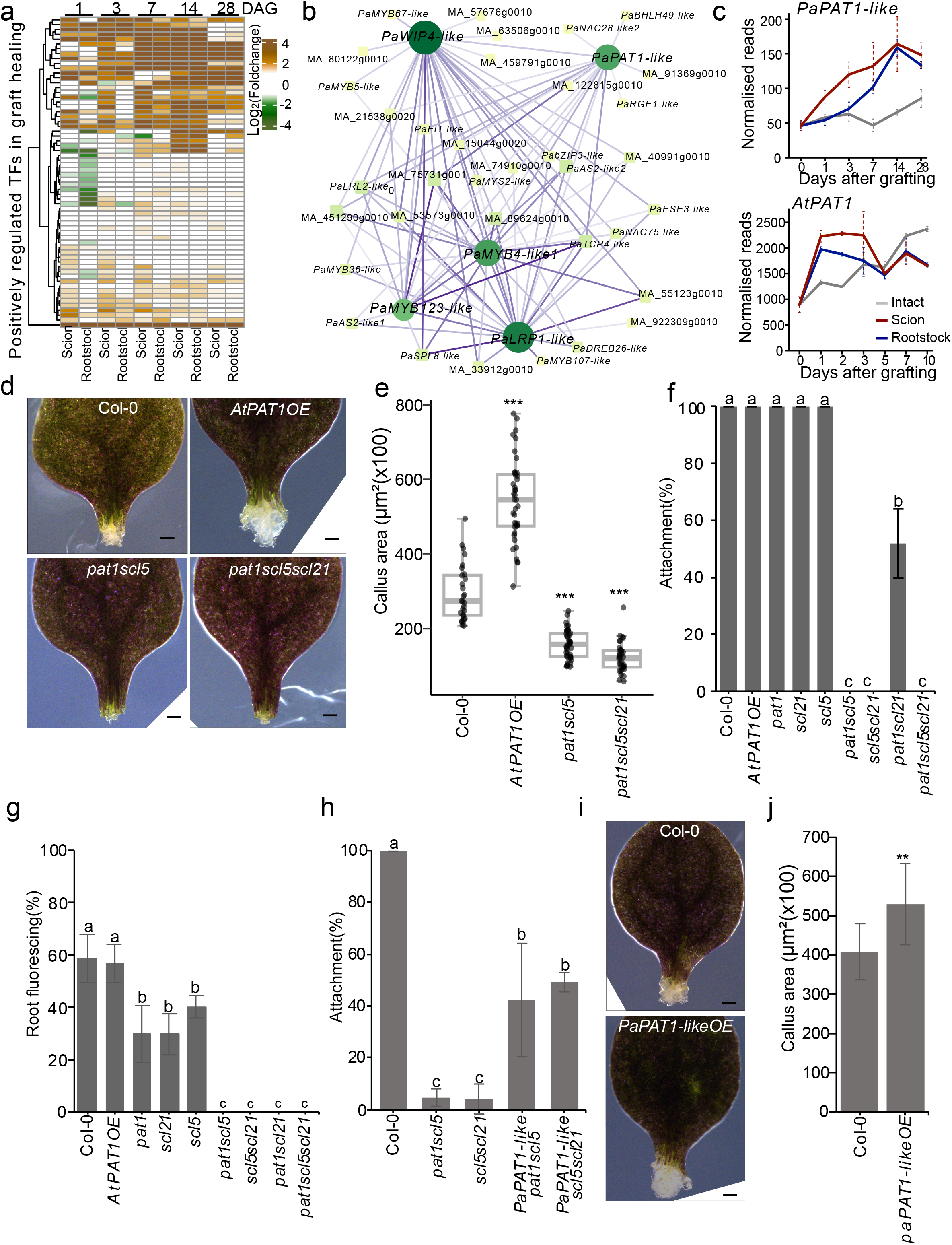
PAT1-related genes promote graft healing. a, Heatmap of positively regulated transcriptional factors during graft healing. b, Regulatory connections of the top five core transcriptional factors based on a gene regulation network of transcription factors activated by grafting. The size and color of nodes indicate the number of edge connections. The edge color indicates the value of correlation. c, *PaPAT1-like* and *AtPAT1* expression during graft formation in *Picea abies* and *Arabidopsis*. d, Images showing callus formation from cut *Arabidopsis* petioles overexpressing *PAT1* (*AtPAT1OE*) or mutant for *pat1scl5* or *pat1scl5scl21*. Scale bars, 100 µm. e, Petiole callus area in wounded petiole explants in Col-0 (n=28), *AtPAT1OE* (n=40), *pat1scl5* (n=41) or *pat1scl5scl21* (n=35). Dots indicate individual samples, *** *p*<0.001. Wilcoxon’s test. (f-g), Attachment and phloem reconnection rates of homografted Col-0, *AtPAT1OE*, *pat1*, *scl5*, *scl21*, *pat1scl5*, *scl5scl21*, *pat1scl21* and *pat1scl5scl21*. Different letters represent *p* < 0.05. One-way ANOVA, with Tukey’s post-hoc test. h, Graft attachment rates of *Picea abies PaPAT1-like* transformed in *pat1scl5* or *scl5scl21* backgrounds. Different letters represent *p* < 0.05. One-way ANOVA, with Tukey’s post-hoc test. (i-j) Images and quantifications of callus formation from cut petioles of Col-0 and *PaPAT1-like* overexpression. One cut petiole from each plant. Scale bars, 100 µm. Student’s t test, ** *p*<0.01. Source data of e, f, g, h and j are provided in the supplementary Table 3.

Our results pointed to an important role for *PAT1* and *ERF115*-related genes in *Arabidopsis* grafting. To investigate this gene family in *P. abies*, we first constructed a phylogenetic tree with *PaPAT1-like* and *Arabidopsis* GRAS family genes. The phylogeny indicated that *PaPAT1-like* was most closely related to *AtPAT1*, *AtSCL5* and *AtSLC21*. An amino acid alignment also showed strong similarity between proteins of these genes (Extended Data Fig. 6a, b and c). To examine if *PaPAT1-like* could perform a similar function as *Arabidopsis PAT1*, we cloned the *P. abies PaPAT1-like* gene to generate an inducible overexpression (OE) line *35S::XVE>>PaPAT1-YFP* in *Arabidopsis* Col-0 (*PaPAT1-likeOE*), *pat1scl5* or *scl5scl21* backgrounds. We found that *PaPAT1-likeOE* could partially rescue the graft attachment defect of *pat1scl5* and *scl5scl21* double mutants (Fig. 6h), and could increase callus formation rates in cut petioles compared to wild-type (Col-0). We also cloned and overexpressed *35S::XVE>>PaERF115-like-YFP* (*PaERF115-likeOE*) in the *Arabidopsis AtPAT1OE* background and found *PaERF115-like*-*AtPAT1* massively increased callus formation (Extended Data Fig. 7). Together, our results indicated that grafting-induced *PAT1* upregulation was shared between *Arabidopsis* and *P. abies* and that this gene has a conserved role in wound healing.

## Discussion

Propagating and combining species through grafting is commonly used worldwide, highlighting the need to establish efficient and practical grafting methods for diverse species ^8, 38, 40^. Here we developed a micrografting method that allows multiple conifer species to be grafted together, expanding the range of compatible graft combinations and providing insights into grafts formation between *Picea*, *Pinus* and *Larix* members that are estimated to have diverged from each other over 100 million years ago ^41^. Previous conifer micrografting methods between mature leaf and young seedling rootstocks had low grafting success rates and seasonal limitations ^7^. In this study, our rapid micrografting approach (50 grafts/hour) provided high success rates in both homografted and inter-species conifer graft combinations. Inter-genus combinations also showed high initial survival frequency though with time, many combinations died or showed stunted growth consistent with delayed incompatibility ^42^.

However, several inter-genus combinations were successful after several years. *P. abies* rootstock were compatible (20-30%) with *P. sylvestris*, *P. contorta* and *L. siberica* scions. Reciprocal grafts with *P. abies* scions had much lower long-term success rates suggesting *P. abies* roots allowed inter-genus graft compatibility whereas scions did not. Previous inter-genus grafts between *Lotus japonicus* and *Medicago truncatula*, or *Solanum lycopersicum* and *Capsicum annuum*, also showed different efficiencies when scion and rootstock genotype were reversed ^34, 43^. The basis for this ‘graft polarity’ is not well known, but *Arabidopsis alf4* mutants deficient in auxin response perturb grafting when in the rootstock but not in the scion, consistent with *Arabidopsis* rootstock being more sensitive to auxin than scions ^17^, and suggesting at least one plausible explanation for differences in grafting efficiencies between scion and rootstock.

Our transcriptome analysis of the *P. abies* graft junction revealed dynamic activation of genes related to cell division, cambial identity, phloem and xylem, similar to what has been previously described at the *Arabidopsis* graft junction (Fig. 3, Extended Data Fig. 3) ^23^. However, there were notable differences. In *Arabidopsis*, phloem reconnection (3-4 DAG) is followed by xylem reconnection (6-7 DAG) ^17^ but our *Picea* transcriptomes showed the homologous genes of *Arabidopsis* xylem and phloem markers activating at similar times. Graft healing also took a longer time in *Picea* and *Pinus.* However, we found multiple similarities regarding the physiology and gene expression changes during grafting, pointing to a conserved process. Our study also identified auxin responsive genes in *P. abies* and found elevated auxin response at the graft junction that correlated with the activation of cell-wall related genes (Fig. 3 and Extended Data Fig. 4). Both auxin and cell-wall related processes promote graft formation in *Arabidopsis*, grape, *Nicotiana* and tomato ^23, 37, 44, 45^, implying that auxin response and cell wall modifications are important for graft formation in both *Arabidopsis* and conifers.

Our gene regulatory network analysis identified several expression hubs including a *PaPAT1- like* homolog of an *Arabidopsis* GRAS transcription, *PAT1*. The PAT1 protein sequence was highly similar (>78% of protein sequence identity in C-terminus) between *Picea* and *Arabidopsis*, in particular the C-terminus found in many GRAS/SCL family members. Our cross-species complementation analysis demonstrated that PaPAT1 was functional and acted similar to AtPAT1 since we observe partial rescue of attachment phenotypes and callus formation. Previous studies have also used genes from tree species to modify *Arabidopsis* function. MADS-box genes *DAL2*, *DAL11*, *DAL12*, and *DAL13* from *P. abies* showed a common function with homologous genes in angiosperms suggesting conservation of genes structure and activity ^46, 47^. A poplar *PeSCL7* can improve salt tolerance of *scl7* mutant in *Arabidopsis* ^48^ whereas several conifer genes including *HBK1*, *SAG1* and *NEEDLY* can functionally substitute for their homologous genes in *Arabidopsis* ^49–51^. Such heterologous techniques are useful tools to understand the function of these genes outside of their endogenous system, particularly with the limitations of CRISPR mutations in trees, however, further work is needed with *P. abies* mutants in *PAT1* to confirm its endogenous role in wound healing and graft formation.

Our work with *P. abies* helped identify *PAT1* as a novel regulator of tissue attachment and graft formation in *Arabidopsis*. We also see evidence that a *PAT1* homolog is induced by wounding in rice (Extended Data Fig. 7)^16^. Monocot grafting succeeds due to the use of embryonic tissues ^16^, and here, the use of young tissues also allowed divergent conifer grafts to succeed. These findings imply that grafting young tissues helps overcome incompatibility in both gymnosperms and angiosperms, and presents a useful tool to extend the range of successful grafting in seed plants.

## Materials and methods

### Plant materials and growth conditions

All conifer seeds (*Picea. abies*: FP-96 Skogsgård was used for grafting and generating graft junction libraries, and FP-518 TreO G7 Söregärde was used for generating hormone treatment libraries; *Pinus. sylvestris*: FP-601 Almnäs; *Pinus. contorta*:FP-704 Lögdö; *Larix* hybrid (*Larix × Marschlinsii*): FP-73 Långtora; *Larix sibirica*: SV309 Lassinmaa) were surface sterilized in 30% hydrogen peroxide solution for 20 min, followed by three rinses in autoclaved water, the sterilized seeds were imbibed overnight in darkness. The seeds were germinated on ¼ Murashige and Skoog medium + 1% sucrose + 1% phytagel, and stored vertically in a dark growth cabinet for 6 days, the seedlings were then moved to a low light condition (covered with a A4 paper) for an additional one day, then seedlings were grown in a plant growth chamber (16/8 h light/dark, ∼110 μmol m^-2^ s^−1^, 22°C, chamber). 10-12 days old seedlings were used for micrografting. The grafted plants were grown in the same plant growth chamber in two months after grafting, followed by growing for one year in greenhouse, then a subset of the plants was moved to grow in a field at the Skogforsk research station, Ekebo, Sweden.

All *Arabidopsis thaliana* mutants used in this study were in the Columbia-0 background. *p35S::ERF115*(*AtERF115OE*), *p35S::PAT1*(*AtPAT1OE*), *pat1*, *erf115*, *scl5*, *scl21*, *erf115pat1*, *pat1scl5*, *scl5scl21*, *pat1scl21*, *erf115scl5*, *pat1scl5scl21* were previously published ^26, 27^. To generate *p35S::XVE>>PaERF115-like-YFP*(*PaERF115-likeOE*)*, p35S::XVE>>PaPAT1-like-YFP*(*PaPAT1-likeOE*), the open reading frame (ORF) sequence without stop of *PaERF115-like* and *PaPAT1-like* were amplified from *P. abies* cDNA and cloned into *pDONR221*. RNA extraction was using CTAB/LiCl method^52^ (see RNA extraction section), cDNA was synthesized using First Strand cDNA Synthesis Kit (K1612, Thermo Fisher Scientific). The primers used for cloning spruce genes were listed in supplementary Table 6. All three segments of the entry vectors carrying promoter of *p35S::XVE* ^53^ followed by entry vector carrying ORF and YFP with terminator, were combined into destination vector *pFR7M34GW* ^54^. The constructs were transformed into GV3101 *Agrobacterium* competent cell for plant transformation. Seeds were surface sterilized for 10 min in 70% ethanol, followed by a rinse with 99.5% ethanol, air-dried and sown on petri dishes containing ½ Murashige and Skoog medium + 0.8 % (w/v) agar. Seeds were stratified at 4 for 48 h, then moved to plant growth chamber (8/16 h, light/dark, 80-100 μmol m^−2^ s^−1^ light intensity) and grown vertically.

### Plant micrografting and measurement of attachment and vascular reconnection

*Arabidopsis thaliana* micrografting and carboxyfluorescein diacetate (CFDA) (VWR International) assays for measuring vascular reconnection were performed according to previously published protocols ^10, 12^. For testing graft attachment, we picked up the cotyledon and root of grafted plants with forceps. Unseparated grafts were counted as attached. For phloem assays, the CFDA was dropped on a cotyledon which was wounded by pressing with forceps. After 1 hour, fluorescence was monitored in the rootstock as an indication of phloem connectivity. Attachment and phloem reconnection were both checked 3 days after grafting.

For conifer micrografting, 10-12 days old plants were excised in the hypocotyl region 0.5 cm below the needles. Scions and rootstocks from different plants were attached tightly together using a 0.8 mm inner diameter silicon collar. To check for the phloem reconnection, we removed all leaves and dipped the cut site into 1mM CFDA solution. Two hours after grafting, we made a hand-section of the rootstocks to observe the CFDA fluorescence. For xylem reconnection assays, we removed the tissues 1 cm below the graft junction, dipping the cut part into 5 μl CFDA for 2 hours, then observed the fluorescence in the needles. To look for xylem differentiation at the junction, we made longitudinal sections at the graft junction, cleared the sections with acidified methanol (Methanol: 37% HCl: H_2_O = 10: 2: 38) ^55, 56^ and incubated at 55-57 for 15-20 min. We then replaced the clearing buffer with 7% NaOH in 60% ethanol and incubated them for 15 min at room temperature, followed by rehydration with 40%, 20% and 10% ethanol, incubating for 15 minutes in each ethanol solution. The rehydrated sections were stained with 0.01% basic fuchsin solution (dissolved in water) for 5 min. The staining was stopped with 70% ethanol for 15 min, and sections were rehydrated with 10% ethanol for 15 min, before adding 50% glycerol and incubating for 30 min. Sections were mounted in 50% glycerol for imaging.

### Imaging

Grafted plants were imaged using a Nikon D5300 camera. The CFDA fluorescence was observed with a Leica M205 FA stereofluorescent microscope fitted with a YFP filter. Basic fuchsin stained samples were imaged with a Zeiss Axioscope A1 microscope or M205 FA stereofluorescent microscope. The higher resolution images of hand-sections were imaged with an LSM-780 confocal microscope. For CFDA imaging, 488 nm excitation and 500-560 nm emission settings were used. 561 nm excitation and 571-690 nm emission were used for imaging basic fuchsin stained samples. All images were analyzed using Fiji^57^.

### Hormone treatment

Two-week-old spruce seedlings were selected for hormonal treatment. Spruce seedlings were treated with either BAP, IAA, or a combination of BAP and IAA hormones. The hormone concentrations were BAP 135 mM, IAA 50 mM, BAP 135 mM + IAA 50 mM. 20 μl of hormone dissolved in 70% ethanol was quickly mixed with a small amount of lanolin. The hormone lanolin paste was then applied to the middle hypocotyl region of the seedlings, approximately 1.5 cm below the needles. The treated area (8-10 mm in length) was wrapped in aluminum foil.

### Construction and analysis of RNA-seq libraries

Graft junction libraries: Samples were collected from both ungrafted (intact) and grafted *P. abies* above and below the graft junction at 0, 1, 3, 7,14 and 28 days after grafting. Approximately 1 mm of tissues from scions or rootstocks or 2 mm from intact plants. Hormone treatment libraries: Samples were collected for analysis (RNA extraction and plastic sectioning) after 5, 10 and 15 days. Approximately 5 mm of the treated hypocotyl was collected from each seedling and the sample was snap frozen with liquid nitrogen.

### RNA extraction

Total RNA was extracted using a modified CTAB/LiCl method^52^. Briefly, frozen tissue was ground into a fine powder. Extraction buffer was prepared (100 mM Tris-HC1 (pH 8), 2% (w/v) CTAB, 30 mM EDTA, 2 M NaCI, 0.05% (w/v) spermidine, 2% (w/v) PVPP, 2% (v/v) 2-mercaptoethanol, proteinase K (10 mg/mL) to a final concentration of 1.5 mg/mL) and warmed for 10 min at 42, then added to the ground frozen tissue and incubated for at 42 for 90 min. Chloroform-isoamyl alcohol (24:1 [v/v]) was added to extract RNA. After vortexing and centrifuging at 15,000 g for 15 min at 4, the top aqueous phase was transferred to a new tube and ¼ vol of 10 M LiC1 was added, allowing overnight precipitation at 4. Samples were centrifuged at 15,000 g for 30 min and supernatant discarded. Finally, the pellet was washed with 2 M LiCl twice and dissolved in RNase-free water. RNA concentration was measured using Qubit 2.0. The RNA Integrity Number (RIN) was analyzed by using an Agilent 2100 Bioanalyzer with RNA 6000 Nano kit. The RINs of all samples were above 8.0.

### Libraries construction

1 ug total RNA per sample was used for RNA-seq library preparation. For grafting junction libraries: mRNA isolation was performed using NEBNext® Poly(A) mRNA Magnetic Isolation Module (NEB #E7490S), followed directly by using NEBNext Ultra Directional RNA Library Prep Kit for Illumina (NEB #E7760S) and NEBNext Multiplex Oligos for Illumina (NEB #E7600S) for library construction. The quality of DNA libraries was checked by with an Agilent 2100 Bioanalyzer DNA High Sensitivity Kit. The hormone treatment libraries were performed by Novogene (UK).

### Bioinformatic analysis

Sequencing was performed on Illumina NovaSeq 6000 system with PE150. RNA-seq analysis was performed as previously described with minor modifications^58^. Briefly, the quality of raw data was accessed using FASTQC (http://www.bioinformatics.babraham.ac.uk/projects/fastqc/). The residual rRNA contamination was removed using SortMeRNA^59^. Data were then filtered using fastp^60^. After both filtering steps, FASTQC was run again to ensure that no technical artifacts were introduced during the pre-processing steps. Filtered reads were aligned to version 1.0 of the Norway spruce genome (retrieved from the PlantGenIE) using STAR^61^. The parameters of RNA-seq data pre-processing followed the previously described guideline^58^. Read counts were quantified by HTSeq^62^ using Norway spruce version 1.0 GFF file (retrieved from the PlantGenIE), with setting -s reverse. Differentially expressed genes (DEGs) were identified using the DESeq2 package in R^63^. In each time point during grafting, certain intact samples were the reference. However, in each time point during growth, day 0 was the reference. Gene with an absolute log2(Fold change) value above 1 and a q-value below 0.05 were considered differentially expressed. The normalised reads obtained from DEseq2 were used for gene expression. The analysis of common patterns of gene expression during the grafting was performed using the Mfuzz package in R^64^. First, differentially expressed genes of grafted tissues (scion and rootstock) from all time points were combined into a list to consider for analysis. Then based on the DEGs list, day 0 intact samples was also included to identify the gene expression common pattern during grafting. GO enrichment analysis of clusters was performed by hypergeometric distribution in R, with an adjusted pvalue < 0.05 and foldchange > 2 as the cutoff to determine significantly enriched GO terms. We downloaded the spruce GO annotation file from the PlantGenIE. The gene co-expression network was conducted using the WGCNA package^65^ in R. Only 524 putative transcriptional factors that differentially expressed during grafting were analyzed. To increase the sample size, the individual replicates were introduced as one sample. Then a module that showed positive regulation during the grafting was considered to discover the hubs of regulatory interactions. Then the interactions were visualised in Cytoscape 3^66^. The orthologs between Norway spruce and Arabidopsis were obtained from pabies_artha.tsv (retrieved from the PlantGenIE). The list of DEG from all comparisons and the orthologs, and TF of the regulatory network are provided in Supplementary Table 7, as well as the genes used for heatmap plots.

### Callus induction

Cotyledons with petioles excised from 10 days old seedlings growing in long day condition were used for callus induction^39^. The cotyledon explants were placed on petiole callus-induction medium (MS medium plates supplemented with 1% sucrose and 0.6% Gelrite) and induced for 7 days under long day conditions. For callus formation without wounding, the F1 generation of *AtERF115OE-AtPAT1OE* seedlings were growing on ½ MS medium for 21 days, and the T2 generation of *PaERF115-likeOE-AtPAT1OE* seedlings were growing on ½ MS medium with 10 µM Dexamethasone (DEX, Sigma-Aldrich) for 7 days, then transferred to ½ MS medium for 14 days.

### Estradiol treatment

For callus induction and grafting assay using *Arabidopsis,* plants growing medium and callus induction medium were both prepared containing 10 μM estradio (Sigma-Aldrich). The grafted plants were place on the filter paper containing water with 10 μM estradiol.

### Phylogenetic analysis and amino acid alignment

The protein sequence of Arabidopsis GRAS family and spruce PaPAT1-like were used for Phylogenetic analysis with MEGA software (version 11) ^67^. The protein sequences were aligned by ClustalW, Construct/Test neighbor-joining tree was used to estimate phylogenetic trees. Bootstrap Replicates number was set 1000. Amino acid alignment was generated with Snapgene software (version 5.1.4.1; www.snapgene.com). The protein similarity and heatmap was generated using TBtools software ^68^.

### Measurement of plant size

For plant height measurements, the shoot length of the above-ground part was measured as the plant height. For the shoot diameter measurements, the shoot diameter of grafted plants was measured at 3 cm above the graft, and the shoot diameter of intact plants was measured at similar site to grafted plants.

### Statistical analysis

Statistical analyses methods were used as indicated in the figure legends. Student’s t test was performed with Excel (version 16.72), Wilcoxon signed-rank test and one-way ANOVA followed Tukey HSD test were performed with R (version 4.0.2).

### Data and code availability

The Gene Expression Omnibus (GEO) accession number for the transcriptomic data reported in this paper is GEO: GSE231633. This study did not generate original code.

## Acknowledgements

We thank Mihaly Czimbalmos for helping maintain trees. We thank Martina Leso for helping draw a conifer carton. MF, AZ and CWM were supported by a Wallenberg Academy Fellowship (2016-0274) and an ERC starting grant (GRASP-805094). VN was supported by a Formas grant (2017-00857) to AC. We thank National Academic Infrastructure for Supercomputing in Sweden (NAISS) and the Swedish National Infrastructure for Computing (SNIC) at UPPMAX, partially funded by the Swedish Research Council through grant agreements no. 2022-22465 and no. 2022-23237, for the computations and data handling.

## Figure Legends

**Extended Data Fig. 1| Graft junction healing dynamics in homografted *Picea abies* and *Pinus contorta*.** Confocal images of xylem anatomy taken 10 days after grafting (DAG), 15 DAG, 20 DAG or 25 DAG. At least three plants were grafted for all combinations and time points. Xylem were stained with basic fuchsin. Scale bars, 100 µm.

**Extended Data Fig. 2| Inter-genus micrografting. (**a, c and e), Representative images of homo-grafted or hetero-grafted *Larix* hybrid, *Larix sibirica, Picea abies*, *Pinus sylvestris* and *Pinus contorta*. Images were taken in November when *Larix* were entering dormancy and loosing their needles. Intact (non-grafted) *Larix* hybrid, *Larix sibirica* and *Picea abies* were used as controls. Scale bars, 10 cm. Inserts show the graft junctions. Scale bars, 1 cm. (b, d and f), Height of intact and 2.5 years old grafted *Larix* hybrid, *Larix sibirica, Picea abies*, *Pinus sylvestris* or *Pinus contorta* (n=1-23 per combination).

**Extended Data Fig. 3| Differentially expressed genes during *Picea abies* grafting.**(a, b), Total differentially or non-differentially expressed genes in grafted *Picea abies* scions, rootstocks or intact tissues. Differential expression was determined by comparing to intact samples at the same time point (a) or by comparing to the day zero intact sample (b). (c, d) Total number of upregulated and downregulated genes in scion, rootstock or intact *Picea abies* tissues. Differential expression was determined by comparing to intact samples at the same time point (c) or by comparing to the day zero intact sample (d). (e, f, g), Clustering analysis of transcriptional dynamics during graft healing. Lines indicate the average of differentially expressed genes in scion or rootstock and the expression profiles for selected *Picea abies* genes plotted over grafting. Dots indicate days after grafting (DAG). The number in the brackets represents the number of genes in the cluster. Cluster number is indicated. *Arabidopsis* homolog expression data are plotted and taken from published transcriptome data^23^.

**Extended Data Fig. 4| Auxin, cytokinin and cell wall related genes expression.** a, Principal component analysis of the gene expression data during hormone treatments. Colors indicate different time points and shapes indicate different treatments including auxin (IAA), cytokinin (BAP) and auxin plus cytokinin (IAA+BAP). b, Total differentially or non-differentially expressed genes in hormone treatments at different time point. c, Average expression of auxin responsive genes during graft healing. d, Average expression of cytokinin responsive genes during graft healing. (e, f) Heatmap showing the fold changes of putative beta-1-4-glucanase genes or putative pectate and pectin lyase genes in graft healing or auxin datasets. (g-k), Expression profiles of *PaLAC1-like* (MA_75861g0010), *PaPME5-like* (MA_10267810g0010), *PaKOR1-like* (MA_69345g0010) and *PaPectin lyase-like* (MA_6752784g0010) in the auxin treatment datasets (g) or grafting datasets (h-k).

**Extended Data Fig. 5| Transcription factor analysis during graft formation.** a, Bar plot showing the distribution of all differentially expressed transcriptional factors during graft healing. X-axis indicates the different families of transcription factors, Y-axis indicates the number of differentially expressed TFs during spruce graft formation. b, Hierarchical cluster tree showing co-expression modules identified using WGCNA. Colors indicate different clusters. c, The association between modules and graft healing. Each row corresponds to a module, labeled with a color as in (b). Each column corresponds to the grafted samples (scion and rootstock) and intact samples. The color of each cell indicates the correlation coefficient between the module and samples. (d-g) Expression profiles for the candidate core transcription factor genes during *Picea abies* grafting including *PaLRP1-like* (MA_2299g0010), *PaWIP4-like* (MA_29238g0010), *PaMYB123-like* (MA_10048467g0010) and *PaMYB4-like* (MA_316475g0010). d and g show the homologs of *PaLPR1-like* and *PaMYB4-like* in the *Arabidopsis* graft formation datasets ^23^. h, Expression profile of *PaERF115-like* (MA_10274g0010) during the *Picea abies* graft healing.

**Extended Data Fig. 6| Phylogenetic analysis and amino acid alignment of *Arabidopsis* PAT1 family proteins and *Picea abies* PaPAT1-like protein.** a, phylogenetic analysis of *Arabidopsis* GRAS transcription family proteins and *Picea abies* PaPAT1-like. Red text indicates PAT1 family proteins. b, Amino acid alignment of the C-terminal of AtPAT1, AtSCL5, AtSCL21 and PaPAT1-like. The consensus sequences are highlighted in yellow. c, Heatmap showing the amino acid similarity in percentages for PAT1 family proteins.

**Extended Data Fig. 7| ERF115 and PAT1 are important for graft healing and regeneration.** a, Graft attachment rates in *Arabidopsis* Col-0, *erf115*, *erf115pat1* and *erf115scl5*. Different letters represent *p* < 0.05. One-way ANOVA, with Tukey’s post-hoc test. b, Phloem reconnection rates in *Arabidopsis* Col-0, *erf115*, *erf115pat1* and *erf115scl5*. One-way ANOVA, with Tukey’s post-hoc test. c, Images showing three weeks old Col-0, *AtPAT1OE*, *AtERF115OE-AtPAT1OE* and *PaERF115-likeOE-AtPAT1OE*. Scale bars for left and middle panels, 1cm. Scale bars for right panels, 1 mm. d, Rice *OsPAT1-like* (Os07g0583600) expression profile in grafted rice adapted from published data^16^

