## Extended Figures 1-7 for "A conserved graft formation process in *Picea abies* and *Arabidopsis thaliana* identifies the PAT gene family as central regulators of wound healing"

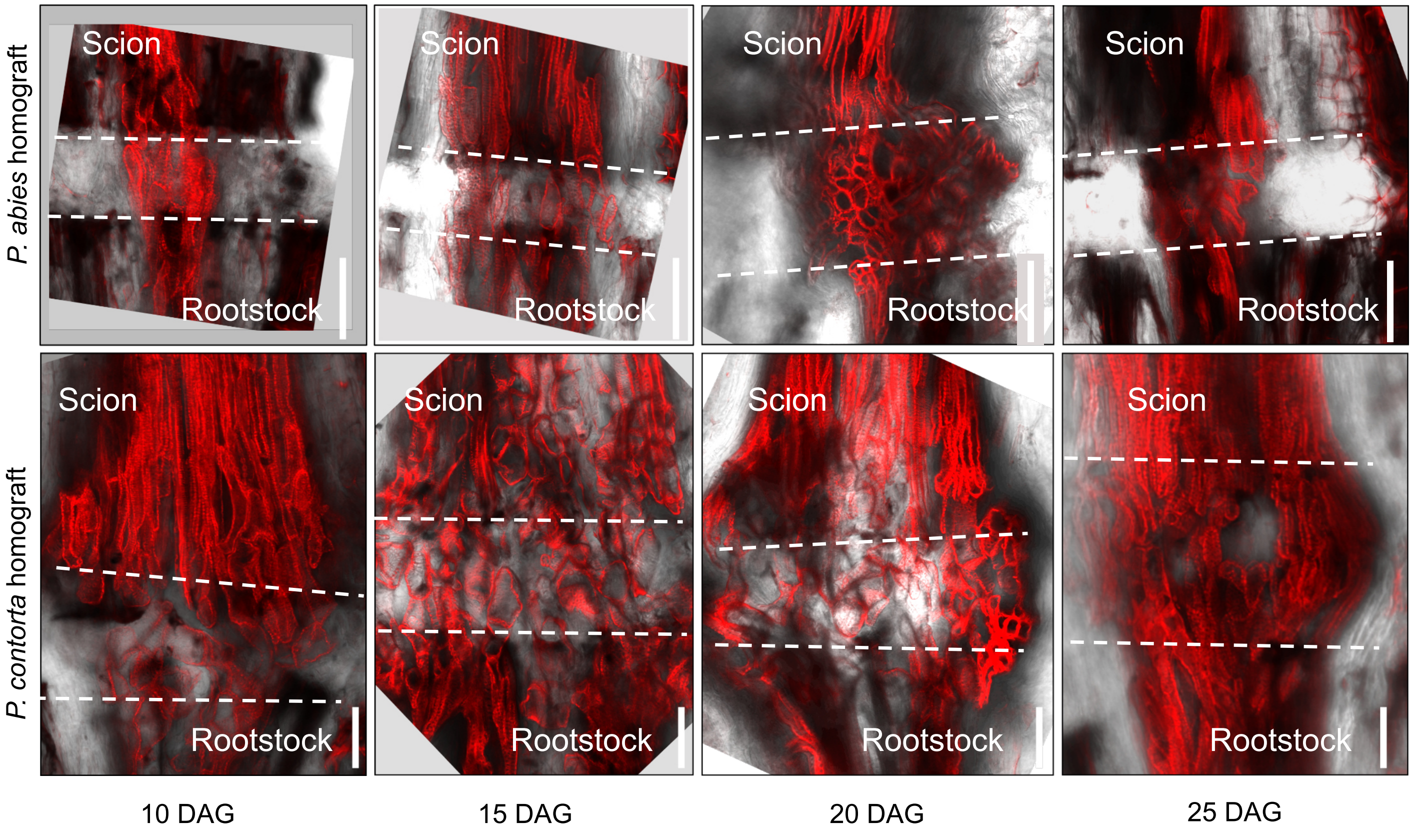


**Extended Data Fig. 1| Graft junction healing dynamics in homografted *Picea abies* and *Pinus contorta*.** Confocal images of xylem anatomy taken 10 days after grafting (DAG), 15 DAG, 20 DAG or 25 DAG. At least three plants were grafted for all combinations and time points. Xylem were stained with basic fuchsin. Scale bars, 100 µm.

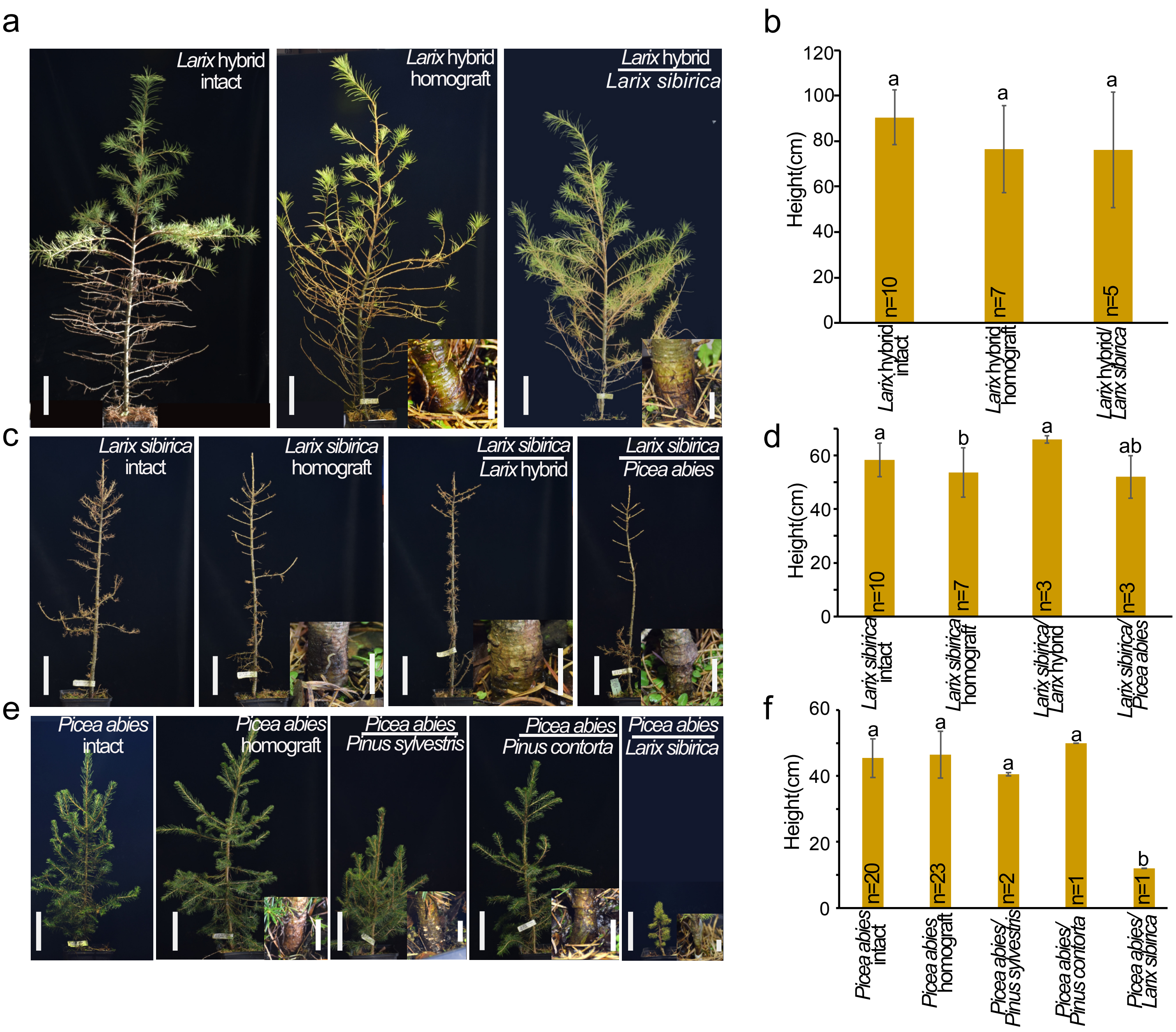


**Extended Data Fig. 2| Inter-genus micrografting. (**a, c and e), Representative images of homo-grafted or hetero-grafted *Larix* hybrid, *Larix sibirica, Picea abies*, *Pinus sylvestris* and *Pinus contorta*. Images were taken in November when *Larix* were entering dormancy and loosing their needles. Intact (non-grafted) *Larix hybrid*, *Larix sibirica* and *Picea abies* were used as controls. Scale bars, 10 cm. Inserts show the graft junctions. Scale bars, 1 cm. (b, d and f), Height of intact and 2.5 years old grafted *Larix hybrid*, *Larix sibirica, Picea abies*, *Pinus sylvestris* or *Pinus contorta* (n=1-23 per combination).

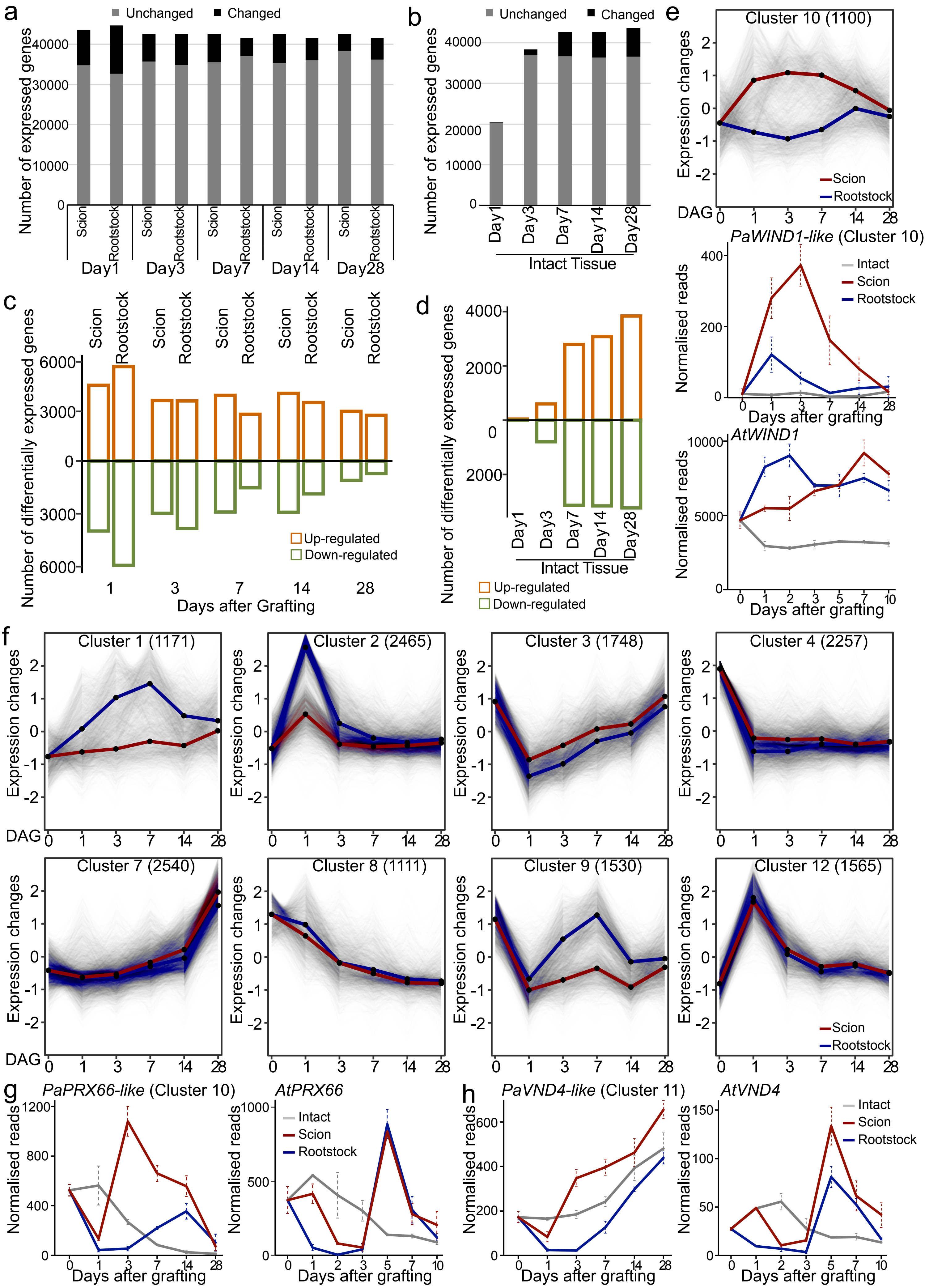

a
Unchanged
Changed
Number of expressed genes
Scion
Rootstock
Day1
Day3
Day7
Day14
Day28
b
Intact Tissue
c
Number of differentially expressed genes
Up-regulated
Down-regulated
Days after Grafting
d
e
Cluster 10 (1100)
Expression changes
DAG
PaWIND1-like (Cluster 10)
Normalised reads
Intact
Days after grafting
AtWIND1
f
Cluster 1 (1171)
Cluster 2 (2465)
Cluster 3 (1748)
Cluster 4 (2257)
Cluster 7 (2540)
Cluster 8 (1111)
Cluster 9 (1530)
Cluster 12 (1565)
g
PaPRX66-like (Cluster 10)
AtPRX66
h
PaVND4-like (Cluster 11)
AtVND4

**Extended Data Fig. 3| Differentially expressed genes during *Picea abies* grafting.**
(a, b), Total differentially or non-differentially expressed genes in grafted *Picea abies* scions, rootstocks or intact tissues. Differential expression was determined by comparing to intact samples at the same time point (a) or by comparing to the day zero intact sample (b). (c, d) Total number of upregulated and downregulated genes in scion, rootstock or intact *Picea abies* tissues. Differential expression was determined by comparing to intact samples at the same time point (c) or by comparing to the day zero intact sample (d). (e, f, g), Clustering analysis of transcriptional dynamics during graft healing. Lines indicate the average of differentially expressed genes in scion or rootstock and the expression profiles for selected *Picea abies* genes plotted over grafting. Dots indicate days after grafting (DAG). The number in the brackets represents the number of genes in the cluster. Cluster number is indicated. *Arabidopsis* homolog expression data are plotted and taken from published transcriptome data[20].

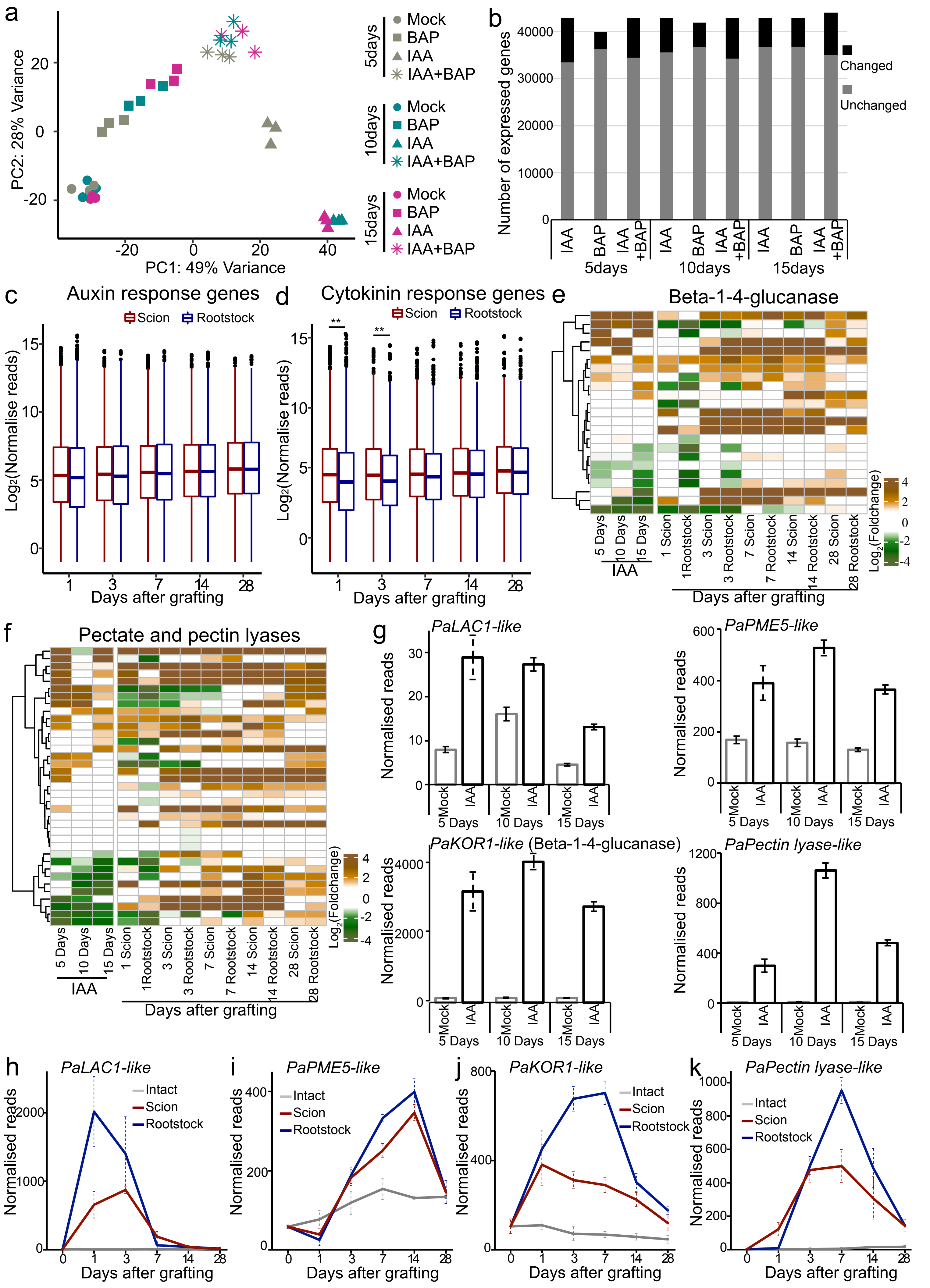

a
PC2: 28% Variance
PC1: 49% Variance
5days
10days
15days
Mock
BAP
IAA
IAA+BAP
b
Number of expressed genes
Changed
Unchanged
IAA
BAP
IAA +BAP
5days
10days
15days
c
Auxin response genes
Scion
Rootstock
Log2(Normalise reads)
Days after grafting
d
Cytokinin response genes
Scion
Rootstock
Log2(Normalise reads)
Days after grafting
e
Beta-1-4-glucanase
5 Days
10 Days
15 Days
IAA
1 Scion
1Rootstock
3 Scion
3 Rootstock
7 Scion
7 Rootstock
14 Scion
14 Rootstock
28 Scion
28 Rootstock
Days after grafting
Log2(Foldchange)
f
Pectate and pectin lyases
5 Days
10 Days
15 Days
IAA
1 Scion
1Rootstock
3 Scion
3 Rootstock
7 Scion
7 Rootstock
14 Scion
14 Rootstock
28 Scion
28 Rootstock
Days after grafting
Log2(Foldchange)
g
PaLAC1-like
PaPME5-like
PaKOR1-like (Beta-1-4-glucanase)
PaPectin lyase-like
Normalised reads
Mock
IAA
5 Days
10 Days
15 Days
h
PaLAC1-like
i
PaPME5-like
j
PaKOR1-like
k
PaPectin lyase-like
Intact
Scion
Rootstock
Normalised reads
Days after grafting

**Extended Data Fig. 4| Auxin, cytokinin and cell wall related genes expression.**

a, Principal component analysis of the gene expression data during hormone treatments. Colors indicate different time points and shapes indicate different treatments including auxin (IAA), cytokinin (BAP) and auxin plus cytokinin (IAA+BAP). b, Total differentially or non-differentially expressed genes in hormone treatments at different time point. c, Average expression of auxin responsive genes during graft healing. d, Average expression of cytokinin responsive genes during graft healing. (e, f) Heatmap showing the fold changes of putative beta-1-4-glucanase genes or putative pectate and pectin lyase genes in graft healing or auxin datasets. (g-k), Expression profiles of *PaLAC1-like* (MA_75861g0010), *PaPME5-like* (MA_10267810g0010), *PaKOR1-like* (MA_69345g0010) and *PaPectin lyase-like* (MA_6752784g0010) in the auxin treatment datasets (g) or grafting datasets (h-k).

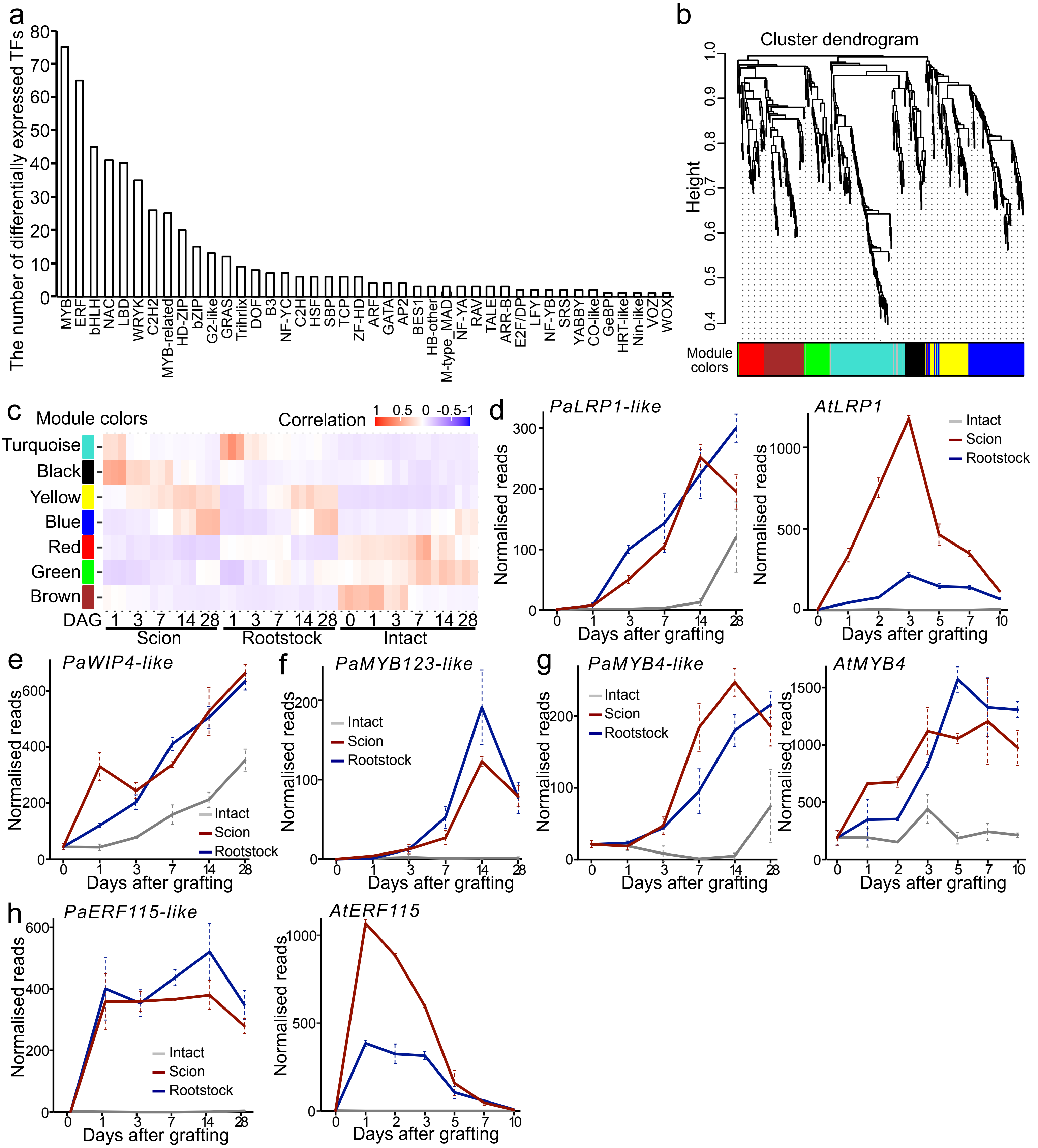


**Extended Data Fig. 5| Transcription factor analysis during graft formation.**

a, Bar plot showing the distribution of all differentially expressed transcriptional factors during graft healing. X-axis indicates the different families of transcription factors, Y-axis indicates the number of differentially expressed TFs during spruce graft formation. b, Hierarchical cluster tree showing co-expression modules identified using WGCNA. Colors indicate different clusters. c, The association between modules and graft healing. Each row corresponds to a module, labeled with a color as in (b). Each column corresponds to the grafted samples (scion and rootstock) and intact samples. The color of each cell indicates the correlation coefficient between the module and samples. (d-g) Expression profiles for the candidate core transcription factor genes during *Picea abies* grafting including *PaLRP1-like* (MA_2299g0010), *PaWIP4-like* (MA_29238g0010), *PaMYB123-like* (MA_10048467g0010) and *PaMYB4-like* (MA_316475g0010). d and g show the homologs of *PaLPR1-like* and *PaMYB4-like* in the *Arabidopsis* graft formation datasets (Melnyk et al., 2018). h, Expression profile of *PaERF115-like* (MA_10274g0010) during the *Picea abies* graft healing.

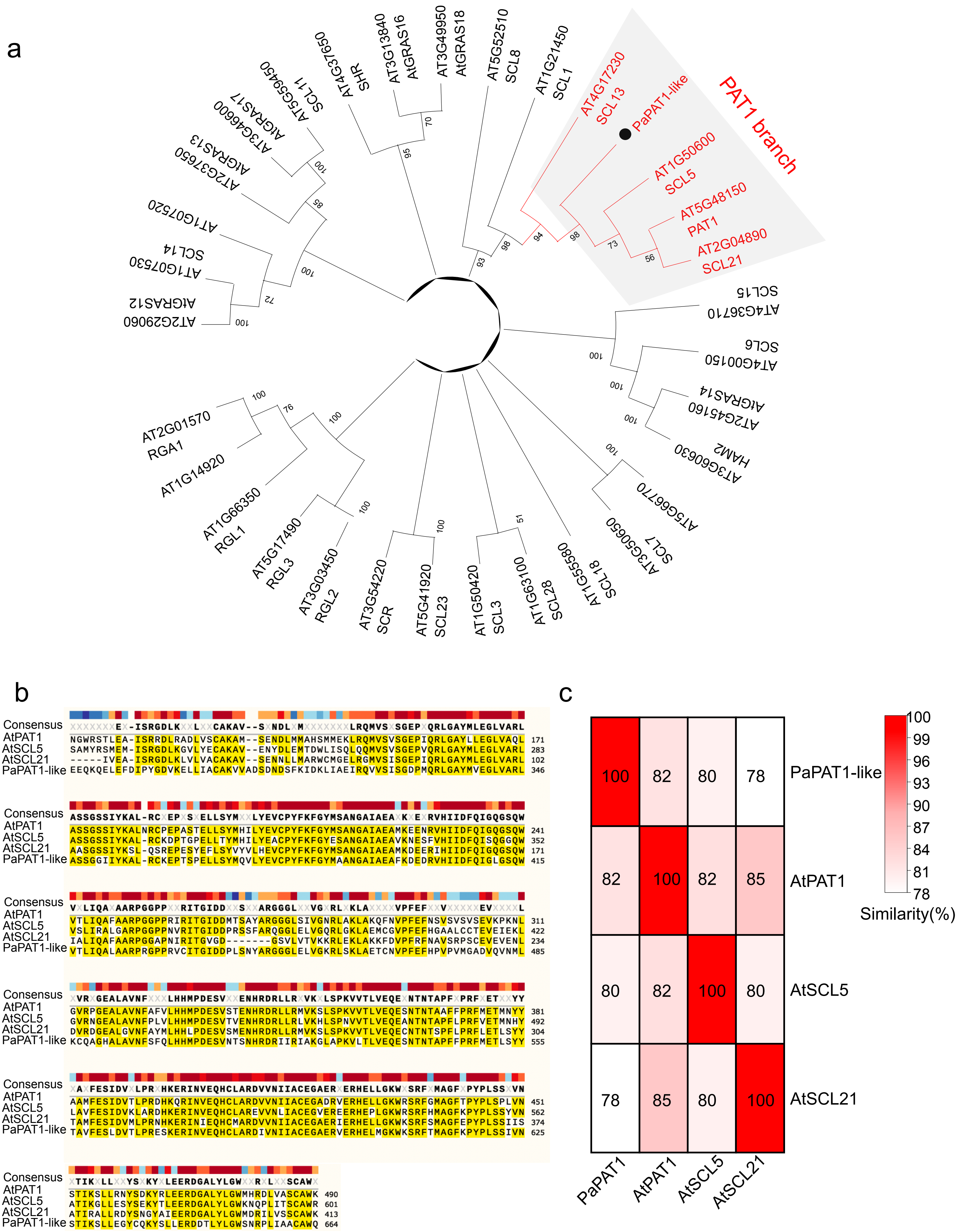


**Extended Data Fig. 6| Phylogenetic analysis and amino acid alignment of *Arabidopsis* PAT1 family proteins and *Picea abies* PaPAT1-like protein.**

a, phylogenetic analysis of *Arabidopsis* GRAS transcription family proteins and *Picea abies* PaPAT1-like. Red text indicates PAT1 family proteins. b, Amino acid alignment of the C-terminal of AtPAT1, AtSCL5, AtSCL21 and PaPAT1-like. The consensus sequences are highlighted in yellow. c, Heatmap showing the amino acid similarity in percentages for PAT1 family proteins.

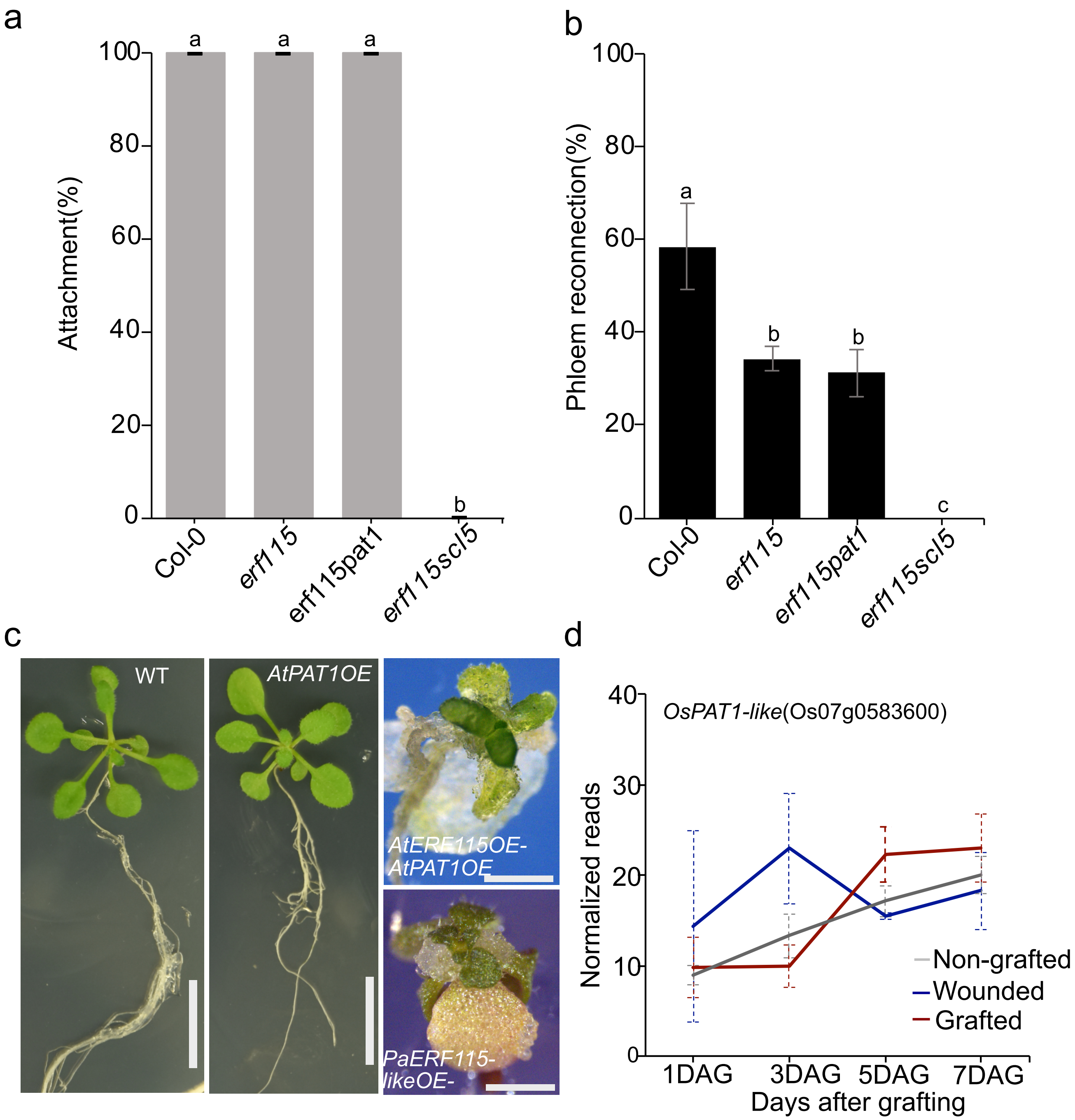


**Extended Data Fig. 7| ERF115 and PAT1 are important for graft healing and regeneration.**
a, Graft attachment rates in *Arabidopsis* Col-0, *erf115*, *erf115pat1* and *erf115scl5*. Different letters represent $p < 0.05$. One-way ANOVA, with Tukey's post-hoc test. b, Phloem reconnection rates in *Arabidopsis* Col-0, *erf115*, *erf115pat1* and *erf115scl5*. One-way ANOVA, with Tukey's post-hoc test. c, Images showing three weeks old Col-0, *AtPAT1OE*, *AtERF115OE-AtPAT1OE* and *PaERF115-likeOE-AtPAT1OE*. Scale bars for left and middle panels, 1cm. Scale bars for right panels, 1 mm. d, Rice *OsPAT1-like* (Os07g0583600) expression profile in grafted rice adapted from published data[13]
